# Analysis of Combinatorial Knockout CRISPR Screens with GRAPE: Genetic interaction Regression Analysis of Pairwise Effects

**DOI:** 10.64898/2026.02.09.704894

**Authors:** Juihsuan Chou, Chenchu Lin, Subin Kim, Junjie Chen, Traver Hart

## Abstract

Genetic interactions (GI) reveal functional relationships for understanding gene function and identifying candidate therapeutic vulnerabilities. Combinatorial CRISPR technologies enable genome-scale GI mapping in mammalian cells, but existing analytical methods lack systematic validation against ground truths. We introduce GRAPE (Genetic interaction Regression Analysis of Pairwise Effects), a computational framework that identifies GIs from pooled CRISPR screens by using linear regression to estimate single-gene phenotypes and detecting deviations from expected double-knockout effects. To enable rigorous benchmarking, we developed Synulator, a pipeline that simulates realistic CRISPR screen data with defined synthetic lethal interactions while preserving experimental noise profiles. In simulated screens, GRAPE achieves greater precision and recall compared to existing methods, particularly for interactions with weaker effect sizes. Applying GRAPE to published combinatorial screens across cell lines and CRISPR platforms demonstrates concordance with original findings while identifying additional high-confidence interactions. GRAPE provides a robust, versatile tool for GI mapping, advancing functional genomics and the systematic discovery of synthetic lethal targets in cancer.

**Teaser:** A regression-based framework and simulations enable accurate detection of GIs from combinatorial CRISPR screens.

## Introduction

Genetic interactions (GIs) occur when the phenotypic impact of a perturbation in one gene (e.g., mutation or deletion) is influenced by the presence or absence of other genes (*1*). These complex relationships are critical for understanding gene function within cellular pathways, providing a comprehensive view that cannot be captured by analyzing single-gene effects alone (*2*).

Genetic interactions can be broadly classified into two categories: synergistic or suppressing interactions. A suppressor interaction occurs when a double mutant exhibits a less severe phenotype than expected based on the combination of individual gene perturbations and typically arises when two genes participate in the same pathway. For example, when two genes encode subunits of the same protein complex, losing either gene disrupts the complex equally, masking the consequence of a second mutation. Conversely, a synergistic interaction is defined by a double mutant exhibiting a more severe phenotype than expected (*3*). The most extreme and clinically relevant form of synergism is synthetic lethality (SL), where individual gene loss is tolerated, but the disruption of both genes results in complete cell death. SL interactions have proven valuable for targeted cancer therapy as demonstrated using PARP inhibitors in BRCA mutated cancers (*4*–*7*).

The emergence of combinatorial multiplex CRISPR technologies has provided the necessary high-throughput capability to measure pairwise gene perturbations at scale (*8*–*24*). In a typical screen, cells are transduced with constructs expressing CRISPR nucleases (e.g., dual Cas9 (*21*–*23*), orthogonal Cas9 (*20*), hybrid Cas9–Cas12a (*18*), enCas12a (*10*–*12, 19*), or CRISPRi/dCas9 (*24*)) and a library of multi-sgRNA constructs targeting genes singly and in pairs, typically using either a one- or two-component lentiviral system. Over several cell doublings, constructs whose targets yield no fitness phenotype – typically the overwhelming majority – maintain their relative abundance, while constructs that reduce cell fitness are progressively depleted from the population. The abundance of each construct is measured by quantitative sequencing, providing read count profiles that reflect relative fitness across perturbations. The theoretical search space for all-by-all pairwise perturbations in the human protein coding gene is massive. With approximately 20,000 protein-coding genes, testing all pairwise combinations would require screening ∼200 million gene pairs, intractable at the current state of the art. Consequently, many research efforts have utilized multiplex screening technology to address more constrained search spaces, focusing on defined functional pathways, such as those within the DNA damage repair (DDR) pathway (*8, 9, 24*), or a curated list of druggable genes to systematically profile combinations within these specific contexts (*23*).

Alongside these advances in genetic perturbation technologies, new bioinformatic methods have been developed for measuring genetic interactions. Indeed, it is common that large-scale screens are published with their own boutique analytical pipelines. Unfortunately, without gold standards for genetic interactions and/or exhaustive validation assays, the relative accuracy of these methods cannot be systematically evaluated. A robust and versatile methodology is required to accurately deconvolute interactions with a range of phenotypic effects and distinguish true interactions from technical noise, and some external framework for comparison should be available. Here we address both areas. We introduce GRAPE: Genetic interaction Regression Analysis of Pairwise Effects, a robust methodology for scoring genetic interactions from multiplex CRISPR screening data. We also introduce a synthetic lethality simulation framework, Synulator, designed to simulate genetic interaction screens while accurately reflecting the noise profile of these experiments. This simulation framework allows for the rigorous and objective comparison of genetic interaction scoring algorithms, thereby providing a valuable resource in functional genomics and cancer biology.

## Results

Genetic interactions are quantitatively defined as the deviation between the observed combined effect of perturbing two genes and an expected value, often defined as the product (or sum, in log space) of their individual effects (Figure 1A). We developed the Genetic interaction Regression Analysis of Pairwise Effects (GRAPE) framework to infer GIs from pooled dual-perturbation CRISPR screens, whether knockout or CRISPRi. By leveraging all measurements that include knockout of a gene, GRAPE estimates single-knockout (SKO) phenotype, and uses these estimates across gene pairs to model expected double knockout (DKO) phenotype. Deviations from this expectation are evaluated as candidate genetic interactions. The approach can accommodate different multiplex CRISPR systems and can incorporate any number of samples or replicates (see Methods). The method is designed for libraries comprising all pairwise combinations of target genes, organized in either symmetric or asymmetric formats (Figure 1B). Given GRAPE’s reliance on measurements across multiple DKO pairs, GRAPE is not suitable for CRISPR libraries targeting only specific gene pairs, e.g. paralogs, (Figure 1B).

**Fig. 1.**
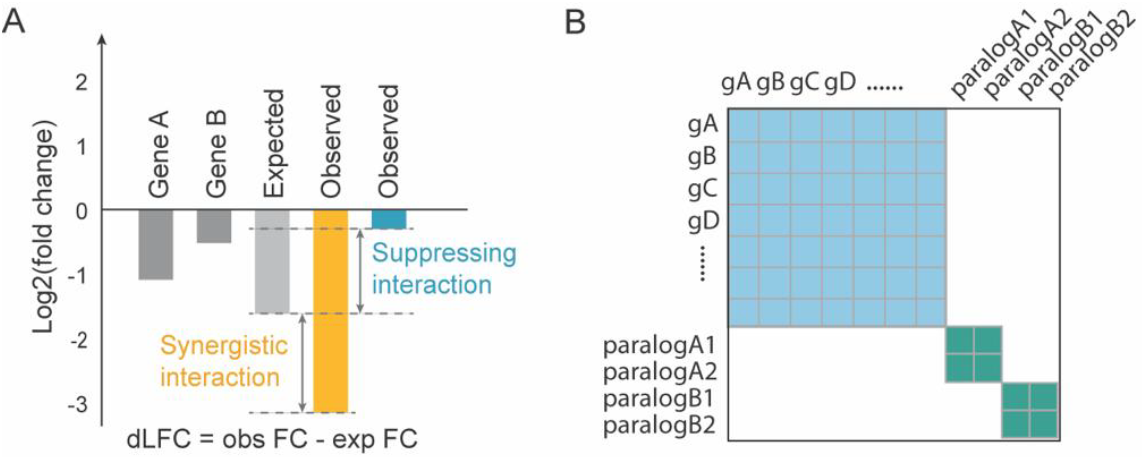
Conceptual framework for measuring genetic interactions in combinatorial CRISPR screens. **(A)** The expected fitness of a dual-gene knockout (KO) is calculated as the sum of the individual single-gene KO log fold change (LFC; y-axis). Delta log fold change (dLFC) is defined as the difference between the observed dual-gene KO LFC and the expected value and serves as a measure of genetic interaction. **(B)** Schematic comparison of CRISPR combinatorial library designs. An all-by-all design targets all possible gene pairs of target genes, whereas a paralog-focused library targets predefined paralog gene pairs.

### GRAPE: Genetic interaction Regression Analysis of Pairwise Effects

A fundamental assumption of large-scale biological experiments is that most observations or perturbations have no effect on the phenotype of interest. GRAPE relies on this assumption by using a linear regression model to estimate single gene knockout phenotypes that best fit the observed double gene knockout phenotypes, measured as log2(fold change) (FC) values, and evaluating deviations from this expectation as candidate genetic interactions (Figure 2A). To demonstrate GRAPE on an experimental dataset, we applied it to the enCas12a CRISPR multiplex screen, as described in Lin et al (*12*), performed in the PC9 cell line using a library targeting all pairwise combinations of 167 DNA damage response genes. GRAPE first normalizes raw read counts and computes the FC for each single and pairwise perturbation. A multiple linear regression model estimates the single gene knockout phenotypes (beta coefficients), expected double knockout phenotype is calculated as the sum of single-gene beta coefficients, and expected FC for each gene pair is compared with observed FC (Figure 2B). To improve robustness, GRAPE offers an optional dynamic range filter that removes the pairs with predicted FCs outside the empirical observed range of the screen (Figure 2B), censoring “more dead than dead” extreme expected fold changes as no-tests. Deviations between observed and expected FCs are defined as raw genetic interaction scores (Figure 2C). To calculate significance of raw GI scores, local variance is estimated using a sliding window of gene pairs with similar expected FC (Figure 2C) and is used to calculate a standardized GI Zscore (Figure 2C-D). In practice, this yields Zgi scores that are normally distributed (Figure 2E), with negative Z-scores indicating synergistic (synthetic lethal) interactions and positive scores indicating suppressing (buffering) interactions. P-values for both negative and positive interactions are calculated from Z-scores, followed by multiple-testing correction using the Benjamini– Hochberg procedure (*25*). We compared Zgi with the Z-transformed delta LFC (ZdLFC) measure, a common method for calculating GI (*22, 26*) and a useful sanity check (Figure 2F). While these two methods are highly correlated (PCC=0.72), Zgi and ZdLFC give substantially different results at stringent cutoffs (Figure 2G). This is largely attributable to the heteroskedasticity filter demonstrated in Figure 2C. Interactions uniquely identified by the ZdLFC metric tended to have more extreme expected FC values (Figure 2H), without correcting for the increased variance in this regime, suggesting that they may be enriched for noise-driven false positives.

**Fig. 2.**
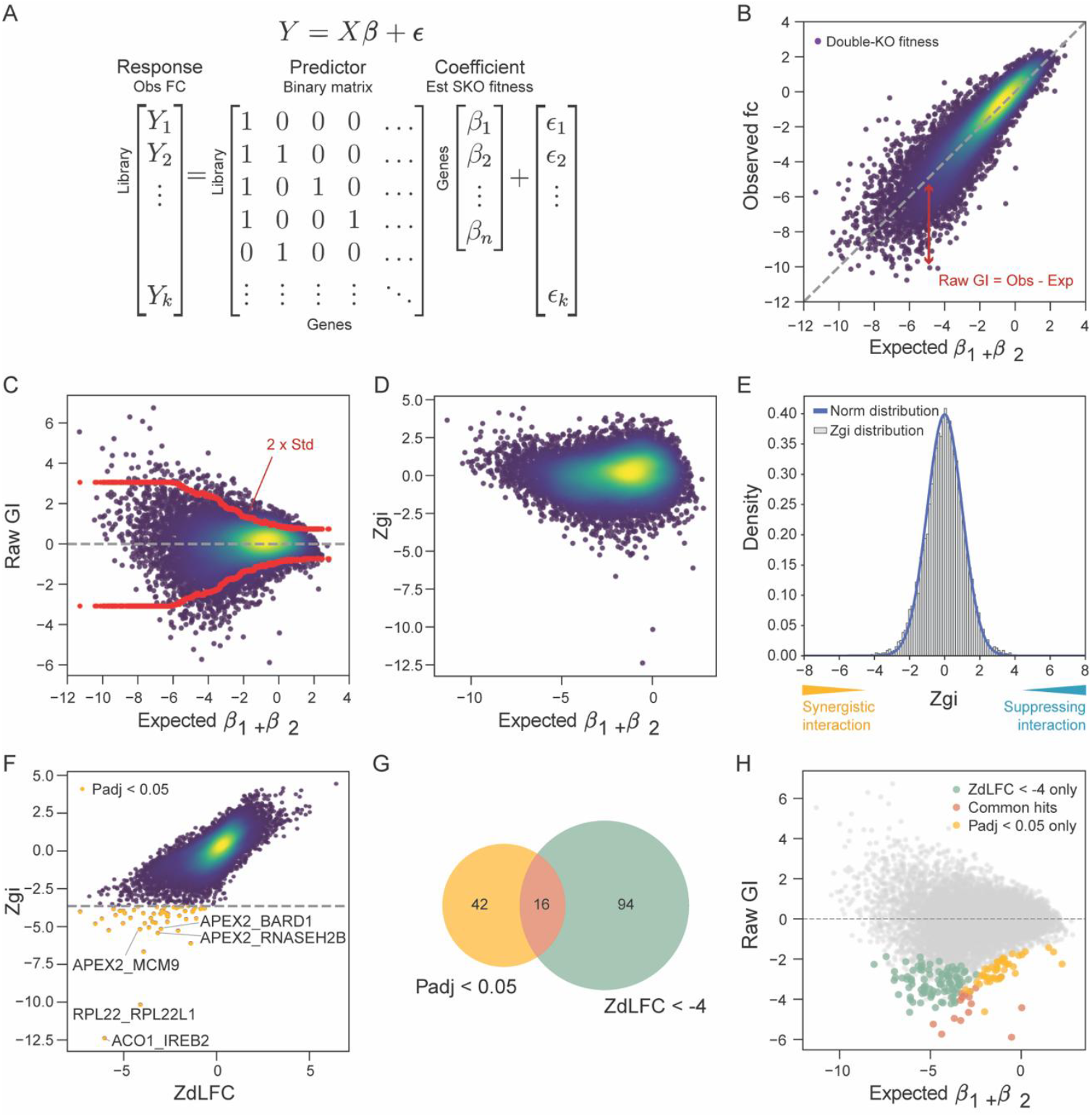
GRAPE method for quantifying genetic interactions. **(A)** Multiple regression framework used by GRAPE. The response vector consists of normalized observed FCs, and the predictor matrix is a binary design matrix indicating the presence or absence of each target gene in a construct. The model estimates single-gene fitness effects (beta coefficients), which are used to calculate expected dual-gene knockout (KO) fitness. **(B)** Scatter plot of observed versus expected FCs for all gene pairs. Deviation from the diagonal represents the raw genetic interaction (GI). **(C)** Raw GI scores plotted against expected FC. Variance increases as expected FC decreases; the red line indicates the local standard deviation used to normalize GI scores and derive GI Zscores (Zgi). **(D)** Scatter plot of Zgi versus expected FC. **(E)** Distribution of Zgi scores, which follows a normal distribution. Negative Zgi values indicate synergistic interactions, whereas positive values indicate suppressing interactions. P values are calculated from Z scores. **(F)** Comparison of Zgi and Z-transformed delta LFC (ZdLFC) scores. Synergistic hits identified using adjusted P < 0.05 are highlighted in yellow. **(G)** Venn diagram showing overlap of top hits identified using Zgi and ZdLFC metrics. **(H)** Scatter plot of Raw GI scores vs expected FC. Hits identified by Zgi (Padj < 0.05; yellow) and by ZdLFC (ZdLFC < -4, green) are highlighted, with common hits shown in pink. Color coding is consistent with panel (G).

Among the top synthetic lethal interactions identified in the DDR library were several biologically well-supported gene pairs (Figure 2F). The paralog pairs *ACO1–IREB2*, which encode iron-responsive element–binding proteins (*27*), and *RPL22–RPL22L1*, ribosomal protein paralogs with compensatory roles in ribosome assembly (*28*), showed strong negative GIs, consistent with paralog buffering and previously reported synthetic lethal enriched in paralogs (*4*). GRAPE also recovered the known synthetic lethal interaction between *APEX2* and *RNASEH2B* (Zgi=-5.45, ZdLFC=-3.16), both of which function in processing aberrant DNA and RNA–DNA hybrids during replication and repair, a relationship reported several times in prior literature (*29*–*31*).

### Synulator: simulation of synthetic lethality for controlled benchmarking of combinatorial CRISPR analysis pipelines

Comparison of the Zgi and dLFC approaches described above highlights a fundamental gap in the analysis of synthetic lethality data in combinatorial CRISPR screens: the absence of a sufficiently large, reliable reference set of synthetic lethal gene pairs against which to benchmark results. To address this gap, we developed a simulation framework, Synulator, to generate realistic synthetic datasets where true interactions with varying effect sizes are known. Synulator simulates CRISPR-based interaction screens by modeling cell proliferation from first principles, using the exponential growth function, *X*_t_ = *X*_0_ · 2^*kt*^, where *X*_0_ represents the initial cell population, *t* is the number of doublings, and *k* is the traditional fitness metric where 1 = wildtype growth and 0 = cell arrest or death. Using this approach, and holding *k* at a fixed value of 1, all wildtype fitness constructs should show a log fold change of exactly zero (see Methods). In contrast, all large-scale CRISPR screens display LFC that are approximately normally distributed around zero (Figure 3A). We account for this by including an error term, *ε*, such that 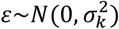, which aggregates biological and technical noise and accurately models the distribution of observed fold changes. A normal fit to the observed fold change distribution has standard deviation equal to *σ*_*k*_ times the number of cell doublings in the screen (Figure 3A).

**Fig. 3.**
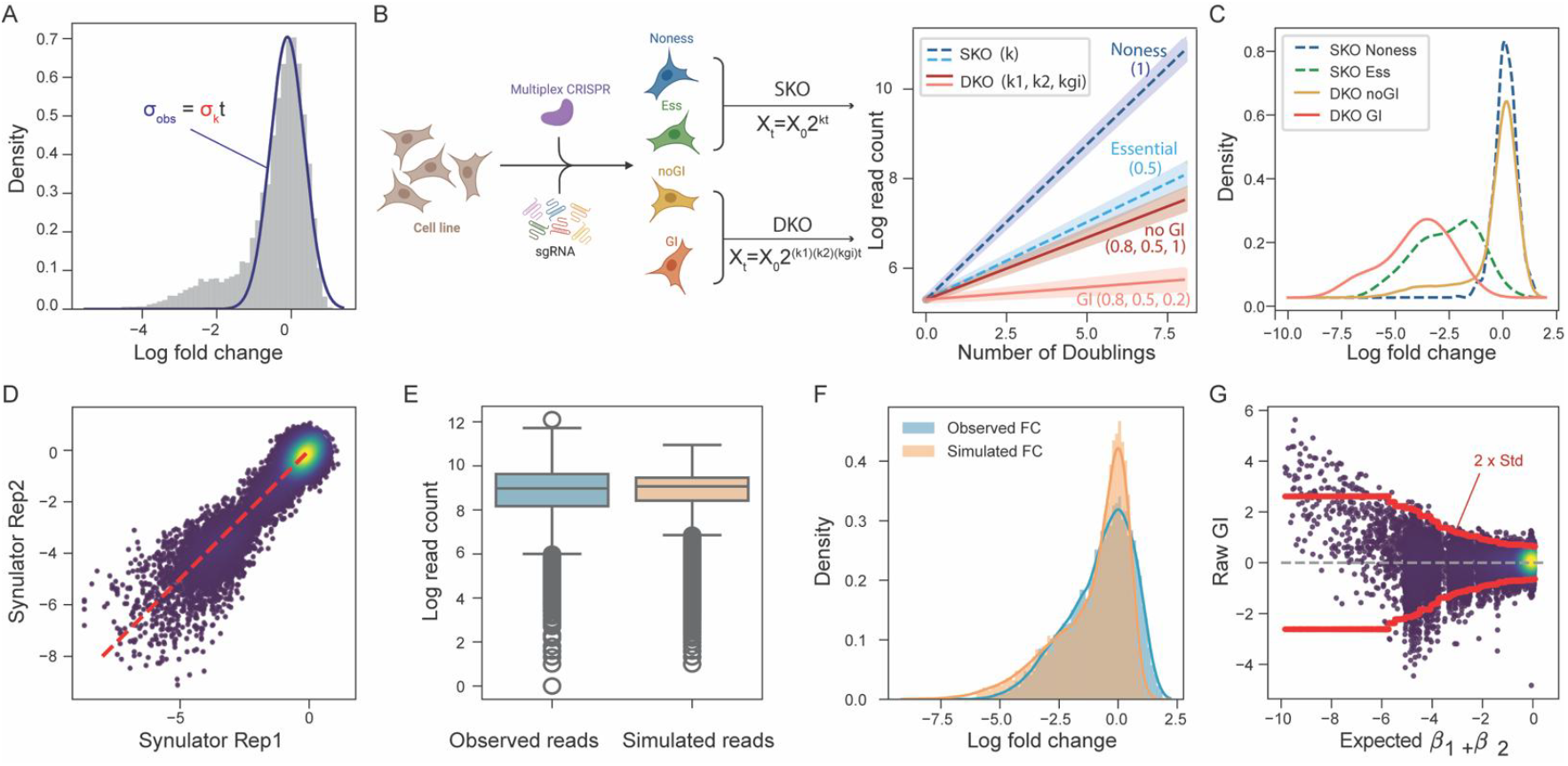
Simulation of combinatorial CRISPR screens using Synulator. **(A)** Empirical FC distribution with nonessential genes centered around zero (dark blue curve). The standard deviation of this distribution *σ*_*obs*_ is used to estimate *σ*_*k*_ the noise term for the fitness parameter *k* in the cell proliferation model *X*_t_ = *X*_0_ · 2^*kt*^. **(B)** Schematic of the Synulator simulation framework. A cell proliferation model is used to generate synthetic CRISPR combinatorial screening data. For single KOs, parameter *k* captures differential growth effects of nonessential versus essential genes. For dual-gene KOs, growth is modeled as *k*_1_ * *k*_2_ * *k*_*gi*_, where *k*_*gi*_ represents the genetic interaction term. Example growth trajectories are shown for nonessential single-gene KO, essential single-gene KO, dual-gene KO without interaction, and dual-gene KO with strong interaction (parameter values indicated). **(C)** FC density distributions for four classes of perturbations: single KO of nonessential genes (dark blue), single KO of essential genes (green), dual KO of gene pairs without genetic interactions (yellow), and dual KO of gene pairs with genetic interactions (red). **(D)** Scatter plot of two independently simulated runs of the same screen, illustrating experimental replicates generated by Synulator. **(E)** Boxplots of log read counts comparing observed data from the PC9 screen with Synulator-generated data. **(F)** FC density distributions comparing observed FCs from the PC9 screen with simulated FCs. **(G)** Application of GRAPE to simulated data, shown as raw genetic interaction scores versus expected FC, analogous to Figure 2C. The red line denotes 2x the local standard deviation.

Under this model, where *k*_*i*_ represents gene *i* knockout fitness, the fitness of combinatorial knockouts is represented by the product *(k*_*i*_*)(k*_*j*_*)(k*_*gi*_), where expected double knockout fitness is the product of *k*_*i*_ and *k*_*j*_, times some genetic interaction factor *k*_*gi*_ (Figure 3B). A *k*_*gi*_ of 1 represents no interaction, with *(k*_*i*_*)(k*_*j*_*)(k*_*gi*_) = *(k*_*i*_*)(k*_*j*_), and *k*_*gi*_ = 0 represents synthetic lethality with a product of zero. Single and double knockouts with and without interactions give rise to variable growth rates (Figure 3B), which translate into expected FC distributions (Figure 3C).

To evaluate how well Synulator can model real experimental data, we parameterized the simulation using the number of genes and guides from our target screen shown in Figure 2 and applied Bayesian optimization to identify the parameter set that best matched the observed read count and LFC distributions. Synulator incorporates negative binomial noise into gene read counts, enabling the simulation to capture sequencing overdispersion and other experimental sources of variability (see Methods). Under the optimized parameter settings (Methods), Synulator closely recapitulates the empirical data (Figure 3D-G). Consistent with empirical CRISPR screens, replicate simulations exhibit increasing dispersion as fold changes decrease (Figure 3D). Notably, when we applied GRAPE to the simulated datasets, the resulting raw GI score distribution (Figure 3G) strongly resembled that derived from the real screen (Figure 2C), demonstrating that Synulator effectively captures the noise structure, overdispersion, and interaction signal present in real CRISPR multiplex experiments.

We used this simulated data, which contains labeled instances of genetic interaction across a range of effect sizes, to compare analytical frameworks developed for interpreting multiplex CRISPR screens. The delta log fold change, dLFC, is calculated as the difference between the observed FC of a gene pair and the expected additive effect, estimated as the sum of the observed single-gene FCs. Horlbeck et al. (*15*) implemented an expanded version of this concept in their dual-CRISPRi study by modeling the expected pairwise phenotypes using a quadratic fitted relationship between single-guide and dual-guide perturbations. GEMINI offers a variational Bayes model for pooled CRISPR–Cas9 double-knockout screens that jointly estimates sample-independent and sample-dependent effects through coordinate ascent variational inference, notionally improving robustness in sparse or noisy datasets (*32*). Orthrus provides a flexible analysis workflow tailored for combinatorial libraries where guide orientation can be explicitly modeled. Its scoring methods compare LFC of targets against controls using either moderated t-testing or Wilcoxon rank-sum testing, depending on the experimental design (*33*). Of the four methods described above, both GEMINI and Orthrus are implemented as generalizable, user-ready software packages. However, continued advances in multiplex CRISPR technology have reduced some design constraints that earlier analytical frameworks were built to address. In particular, the In4mer system leverages constructs encoding up to four gRNAs, enabling two guides per gene in dual target designs and thereby increasing targeting efficiency while maintaining a compact library (*11*). This architecture allows measurements to be obtained with as few as two arrays per gene or gene pair, significantly reducing experimental footprint. As a result, we did not include Orthrus in this simulation benchmark, as its statistical testing framework requires more reagents per gene or gene pair.

### GRAPE demonstrates improved performance to existing methods using simulated genetic interaction data

We compared the performance of GRAPE with existing methods using simulated datasets. We generated dual-perturbation viability screens using Synulator, which provides ground-truth GI gene pairs and therefore enables direct benchmarking of model accuracy. We applied three analytical approaches—the dLFC method, GEMINI, and GRAPE—and assessed their ability to recover labeled genetic interactions. GRAPE achieved the highest PR performance (AUC = 0.869), outperforming both dLFC (AUC = 0.785) and GEMINI (AUC = 0.408) (Figure 4A). We noted that the GI modeled in the simulation showed a range of effect sizes (*K*_*gi*_, Figure 4B), and that performance of both GRAPE and dLFC was highest against high effect size GI and dropped as low effect size GI were included (Figure 4C). GEMINI exhibited relatively poor performance across the board, reliably identifying only interactions with the strongest effect sizes and suffering from a high false-negative rate for all classes of interaction (Figure 4C-D).

**Fig. 4.**
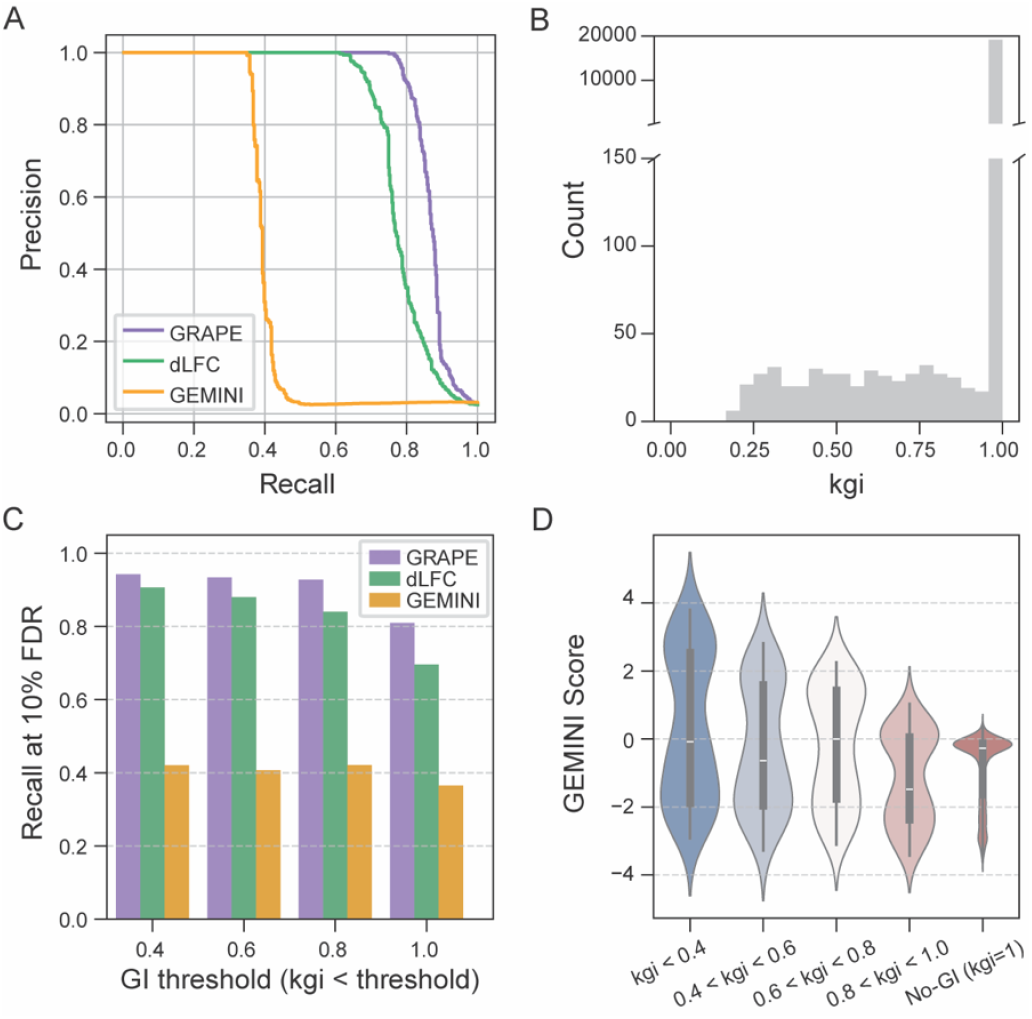
Benchmarking genetic interaction methods using simulated data. **(A)** Precision–recall curves comparing the performance of GRAPE, delta log fold change (dLFC), and GEMINI using Synulator-generated data. **(B)** Distribution of simulated genetic interaction strengths *k*_*gi*_. Most gene pairs have no genetic interactions *k*_*gi* = 1_, whereas decreasing *k*_*gi*_ values correspond to increasingly stronger genetic interactions. **(C)** Recall at 10% false discovery rate (90% precision) for the three methods across different genetic interaction calling thresholds.**(D)** Violin plot showing the distribution of GEMINI strong stratified by genetic interaction strength *k*_*gi*_ bins.

In addition to better performance against benchmarks, GRAPE is significantly easier to use. GEMINI required substantially longer runtime due to iterative variational optimization (approximately 5-7 minutes per run) and, in our experience, often needed multiple attempts to achieve convergence with appropriate parameter settings. In contrast, GRAPE executed the full analysis pipeline—including normalization and QC—typically in less than 10 seconds, whether from the command line or imported as a module into a Python notebook.

### Application of GRAPE to published combinatorial CRISPR screens

Beyond applying GRAPE to our In4mer-based screen, we evaluated its generalizability using three published combinatorial CRISPR datasets that span distinct endonuclease systems and library architectures: the enCas12a dual-guide screens from Lenoir et al. (*10*), the Cas9 dual-promoter system from Fong et al. (*23*), and the dCas9–KRAB CRISPRi screens from Fielden et al. (*24*) These datasets differ in design: Lenoir et al. and Fong et al. performed asymmetric screens (8×92 and 67×176, respectively), whereas Fielden et al. implemented a large all-by-all (548×548) combinatorial library in a single cell line. Applying GRAPE across these datasets demonstrates that the pipeline adapts readily to both symmetric and asymmetric library formats, as well as diverse CRISPR systems. For asymmetric libraries, GRAPE requires only to combine the query and target gene lists into one file and specify via the *--target-gene-file* argument, after which the pipeline conforms to the underlying design.

Figures 5A–5D illustrate GRAPE’s application to the Lenoir et al. MOLM13 dataset. The predicted additive fitness values derived from GRAPE’s regression model mostly track the observed pairwise knockout FCs, resulting in an expected linear relationship for the majority of gene pairs that do not interact (Figure 5A). Top GRAPE hits include *ACACA_GPAT4* and *SREFBF1_SREFBF2*, where *SREFBF1_SREFBF2* is a paralog pair and previously reported synthetic interaction (*34*). In MOLM13 cells Lenoir et al. reported synthetic interactions between *ACACA* and *GPAT4*, consistent with a rewired lipid-metabolism network and increased sensitivity to saturated fatty-acid stress. Because both genes contribute to the same lipid-biosynthetic axis, dual perturbation produces a strong, reproducible fitness phenotype in backgrounds that depend on that pathway (*10, 34*). Another interaction consistently identified by both Lenoir et al. and GRAPE was *ACACA* and *SQLE* (Figure 5D). Although this gene pair has not been validated as a synthetic lethal interaction, both genes encode enzymes in major lipid biosynthesis pathways—fatty acid synthesis (*ACACA*) and cholesterol biosynthesis (*SQLE*). Simultaneous disruption of these pathways might create metabolic stress beyond what either single perturbation causes, providing a potential mechanistic basis for the observed synergistic genetic interaction in MOLM13 cells.

**Fig. 5.**
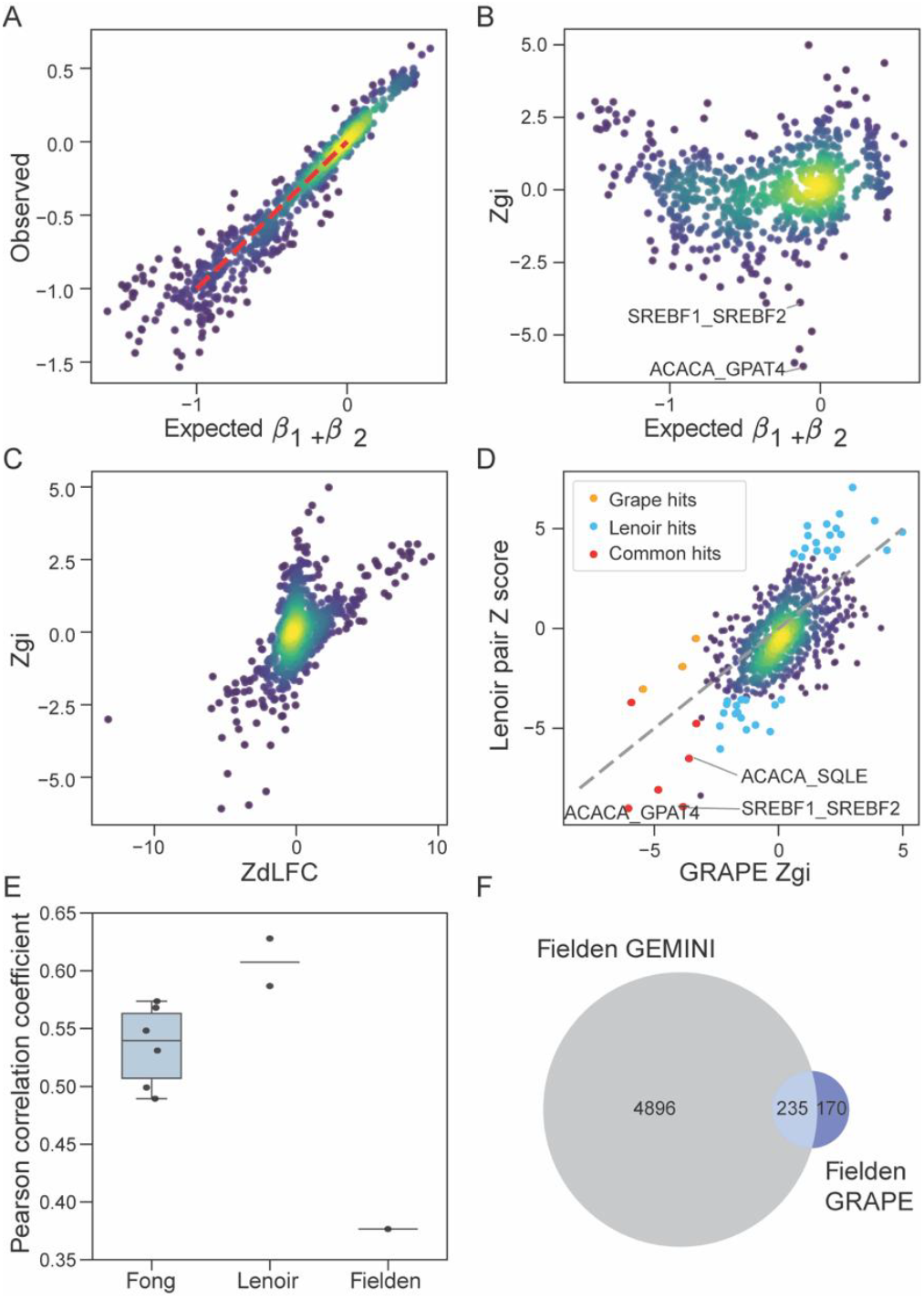
Application of GRAPE to published combinatorial CRISPR screens. **(A)** GRAPE analysis of a combinatorial CRISPR screen from Lenoir et al. in the MOLM13 cell line. Scatter plot shows observed versus expected FCs. **(B)** Scatter plot of GRAPE-derived Zgi versus expected FC. **(C)** Comparison of Zgi and Z-transformed dLFC (ZdLFC) scores. **(D)** Scatter plot comparing genetic interaction scores reported by Lenoir et al. with GRAPE-calculated Zgi values derived from the same raw data. GRAPE-only hits are highlighted in yellow (Padj<0.05), Lenoir-only hits (Padj < 0.01) in blue, and shared hits in red. **(E)** Pearson correlation coefficients between published genetic interaction scores and GRAPE-derived Z_GI values across multiple studies and cell lines. Fong et al. include six cell lines (A549=0.57, CAL27=0.49, CAL33=0.55, MCF10A=0.53, MCF7=0.50, and MDA-MB-231=0.57); Lenoir et al. include MOLM13=0.63 and NOMO1=0.59; Fielden et al. include RPE1=0.38. **(F)** Venn diagram showing overlap among synthetic-lethal hits identified by Fielden et al. using GEMINI (score < – 1), and the same Fielden dataset processed with GRAPE (adjusted P < 0.05).

We also applied GRAPE to the published datasets from Fong et al. (*23*) and Fielden et al.(*24*) (Supplementary Figure 1-2, Supplementary Data S2), analyzing each cell line independently. When comparing the reported GI scores from each study with the GRAPE-derived GI Z-scores computed directly from their raw read counts, we observed Pearson correlation coefficients (PCCs) of approximately 0.5–0.6 for most cell lines, with the Fielden et al. RPE1 screen yielding the lowest concordance (PCC = 0.38) (Figure 5E). This discordance is further illustrated in Figure 5F, which compares hits overlap between GEMINI-based interactions reported by Fielden et al. and GRAPE-derived interactions computed from the same dataset. The limited overlap among these sets shows how different analytical approaches applied to the same underlying data can lead to divergent results. In the absence of an external gold standard, determining which algorithm more accurately discriminates genetic interactions is difficult. Moreover, although Fielden et al. screened a large combinatorial library, the two interacting gene pairs they rigorously validated provide too little ground truth for systematic benchmarking.

## Discussion

In this work, we present GRAPE, a regression-based framework for calling genetic interactions from pooled combinatorial CRISPR screens. By leveraging the shared structure of pairwise perturbations, GRAPE infers single-gene fitness effects directly from dual-knockout measurements and models expected additive fitness for each gene pair. Our analyses using both experimental and simulated datasets show that GRAPE is accurate, computationally lightweight, and adaptable across CRISPR modalities and library architectures.

Published genetic interaction screens often target widely divergent gene pairs and yield divergent hits even in the small areas where their assays overlap. Unfortunately, an even smaller number of gene pairs from these studies are experimentally validated, and each publication uses different hit calling criteria, making cross-study comparisons inconsistent. These limitations make it difficult to objectively evaluate informatic methods, new or old. Together, these observations directly motivate the need for Synulator, which generates realistic synthetic datasets with known ground-truth synthetic interactions, enabling rigorous benchmarking. Synulator generates synthetic interaction data with realistic fitness variation, sequencing overdispersion, and tunable SL interaction strength.

Using Synulator as ground truth, GRAPE outperformed both the standard dLFC method and the variational-Bayes–based GEMINI model, achieving higher precision–recall performance with substantially shorter runtime. Importantly, GRAPE remained robust even as subtle interactions were introduced into the dataset, although, unsurprisingly, detection becomes more challenging at small effect sizes. This reflects an inherent limitation of any genetic interaction calling framework: weaker GIs are often indistinguishable from biological and technical noise in a single screen. However, in principle, replicate screens or multiple perturbation contexts can amplify weak signals. For example, Aregger et al. (*34*) and Lin et al. (*12*) demonstrated that repeated screens in matched knockout backgrounds substantially increase sensitivity for low-effect-size interactions, and recent work from (*35*) relies on multiple replicate screens to confidently identify interactions.

Despite its advantages, GRAPE has several limitations. First, its dependence on adequate library coverage: GRAPE requires that each target gene appears multiple times in the DKO design. Therefore, it is not suitable for paralog-focused screens or gene families where many genes appear only once. Low-effect-size GIs (i.e., *k*_*gi*_ values ∼1) may be indistinguishable from noise in single screens. However, replicate screens or multi-context screens can improve signal-to-noise ratios. Finally, GRAPE assumes broadly linear relationships between observed and expected FCs, consistent with standard GI modeling; however, highly noisy datasets or those with strong nonlinear behavior may require extended modeling strategies.

## Materials and Methods

### GRAPE

#### Calculating fold change and normalization

GRAPE analysis begins with processing the raw read count data obtained from next-generation sequencing and mapping for each sample. A pseudocount of one is added to all counts by default, then normalized by the total read count per sample. Guide-level log_2_ fold changes (FC) are computed as the ratio of read counts at the endpoint divided by T_0_. To obtain gene-level FCs, guide-level values are averaged across all guides targeting the same gene, and the mean across replicates are used for subsequent analyses.

Two normalization options are available for preparing FC input to the regression model. In the default mode-centering approach, FCs are adjusted so that the overall distribution centers on zero, under the assumption that most genes and gene pairs in the library are nonessential and lack strong phenotypes. This ensures that the highest-density region of the FC distribution corresponds to no effect. Alternatively, normalization can be based on a user-specified list of nonessential reference genes provided through the *--nonessential-gene-file parameter*. In this case, GRAPE centers the FCs using the median value of the reference set, ensuring that the nonessential control genes define a baseline centered on zero.

#### Regression modeling and genetic interaction scoring

To estimate single-gene fitness effects and compute genetic interaction (GI) scores, we implemented a linear regression model in the GRAPE pipeline. A binary predictor matrix *A*_*i,j*_ was generated, where each row (*i*) represents a perturbation array in the library and each column (*j*) corresponds to a unique target gene present in the array. *A*_*i,j*_ = 1 if array *i* contains a targeting gene *j*, and 0 otherwise. Rows corresponding to perturbations not targeting any genes in the analysis set, and columns corresponding to genes without targeting arrays, were excluded.

Let y denote the mode-centered observed gene-level log_2_ FC vector, with one value per perturbation array. The matrix *A* and response vector *y* together form the design and response data for regression fitting. We fit a linear model to estimate single-gene knockout effects:

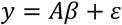

where *β* is a vector of regression coefficients representing the inferred single-gene fitness effects, and *ε* is the residual error. The model was fit without an intercept term (i.e., assuming the mean-centered input FCs were already normalized to zero). Regression was performed using ordinary least squares (OLS). Using the fitted coefficient 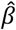, the predicted additive FC for each perturbation was computed as

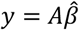

For each gene pair (g1, g2), the expected FC is 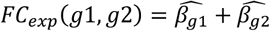, and the observed value from the experiment is *FC*_*obs*_(*g*1,*g*2). The deviation from the additive model defines the raw GI score:

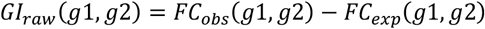

To facilitate comparison with the standard definition of genetic interaction scoring, the observed single-gene FCs were retrieved, and the dLFC was calculated and recorded for each gene pair.

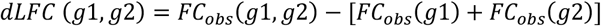

#### Dynamic range filtering

Following regression, gene pairs with predicted FC values outside the assay’s measurable dynamic range were removed prior to normalization. Specifically, we excluded gene pairs whose predicted FC (*FC*_*exp*_) was smaller than the minimum observed FC 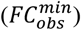 across all perturbations. After filtering, the regression model was refit using the remaining FCs to obtain updated single-gene coefficients and predicted FC.

#### Empirical Bayes Z-score normalization

Because the variance of the raw genetic interaction scores increases as predicted fitness effects become more negative, we applied a local variance normalization approach (*36*). Pairs were ordered by their predicted FC (*FC*_*exp*_), and a local standard deviation (*σ*_*local*_ was estimated within a symmetric window size 2*w* (default w=500) centered on each gene pair. Outliers within each window were excluded using the interquartile range (IQR) rule. Finally, the standardized GI-Zscore was obtained by dividing the raw GI by the local variance estimate:

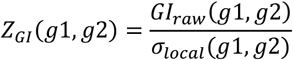

If *w*=0, a single global standard deviation across all pairs was used instead. The optimal half-window size (*w*) should be adjusted according to the total number of gene pairs in the library. In general, *w* should be large enough to include sufficient data points for a stable variance estimate but small enough to preserve local variation across the predicted FC range. As a practical guideline, *w* can be set to approximately 1-2% of the total number of gene pairs in the library. For example, in a library containing ∼35,000 gene pairs, a half-window size of 500 (≈1.4% of the library) provides a good balance between smoothing and resolution. Smaller window sizes (e.g. *w=250*) may better capture local variance, while larger windows (e.g., *w=1000*) may be preferable for noisier screens to stabilize the variance estimate. Users can evaluate the smoothness of the resulting variance curve (Figure 2C) when adjusting *w*. Finally, an optional monotonic variance filter *(--monotone-filter = True*) can be applied to enforce nondecreasing variance estimates along the ordered FC range, preventing spurious dips in variance.

#### Statistical significance

For each gene pair, one-sided p-values were computed under a standard normal distribution to assess either synthetic (negative GI) or suppressive (positive GI) interactions:

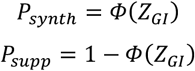

where *Φ*(·) denotes the cumulative distribution function of the standard normal distribution. False discovery rates (FDR) were then estimated using the Benjamini-Hochberg correction, yielding adjusted p-values for calling statistically significant synergistic and suppressive interactions, respectively.

### SYNULATOR

#### Cell Proliferation Model

Cell populations were modeled as an exponential growth process representing the cells transduced with the same gRNA. For each guide, the expected number of cells after *t* doublings was expressed as:

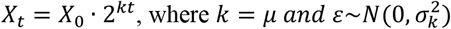

where *X*_0_ is the initial cell count after transduction, *μ* denotes the gene-specific proliferation rate, 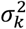 captures the experimental variability empirically observed in fitness screens, and *t* represents the number of doublings.

#### Gene-Level Fitness Model

Gene fitness parameters were defined to capture both single-gene and dual-gene knockout (KO) effects. For gene *i* and *j*, the proliferation rate *Fi,j* was specified as

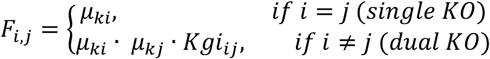

where *μ*_*ki*_ denotes the proliferation rate of individual gene *i*. For dual-gene KO, the proliferation rate is modeled as the product of the two individual gene rates, scaled by a gene-gene interaction term *Kgi*_*ij*_.

Wild type (nonessential) genes were assigned a baseline proliferation rate of 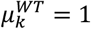. Fitness (essential) genes had their proliferation rate sampled from a uniform distribution within a user-defined range, where the minimum represents strongest guide efficiency:

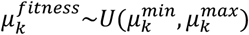

For the genetic interaction parameter *Kgi*_*ij*_, a scaling factor to adjust the genetic interaction frequency were multiplied when both genes are wild type genes. Otherwise, *Kgi*_*ij*_ values were introduced with probability *p*_*GI*_. When present, they were sampled from a uniform distribution:

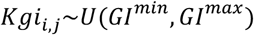

The scaling factor and genetic interaction frequency allows Synulator to capture variable interaction frequencies across gene pairs while maintaining a biologically realistic proportion of interacting genes.

#### Guide-Level Fitness Model

Each gene or gene pair was targeted by *N* gRNAs. To account for both the variation in gene-level variability and guide editing efficiency, guide-level proliferation rate 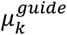 was modeled hierarchically as:

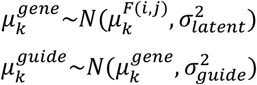

Where 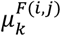 denotes the gene or gene pair specific proliferation rate derived from gene-level fitness model. The variance components were defined such that, after averaging over N guides, the total variance of gene-level means equals the intended fitness noise variance 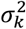 :

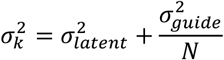

To ensure the latent gene-level variance remains nonnegative, the maximum allowable guide-level variance was used:

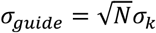

This guarantees that the observed variability across averaged guide measurements matches the desired experimental variance 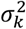, while adding guide-level dispersion.

#### Modeling Experimental and Sequencing Noise

To model the overdispersion typical of sequencing data and experimental noise, observed read counts were simulated using a negative binomial distribution. The mean parameter *n* was scaled by a dispersion parameter *p*:

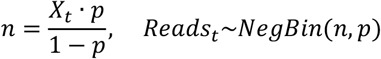

where *Reads*_*t*_ denotes the observed read count for a given guide at *t* number of doublings. This noise model reproduces the noise profile consistent with experimental CRISPR screening data.

#### Calculation of Log Fold Change

Guide-level effects were quantified as log2 fold change (FC) in relative abundance between the initial and final cell populations. The FC for each guide was computed as:

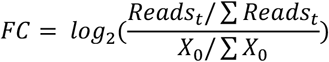

### Running GEMINI using simulated combinatorial CRISPR data

We generated synthetic multiplex CRISPR data using Synulator. Simulations were performed with the following parameters: *num_total_genes = 200, num_fitness_genes = 50, num_guides = 2, median_read_depth = 1000*, and *time = 6*; all remaining parameters were set to default values. Three independent datasets were produced to serve as biological replicates.

Simulated read-count matrices were reformatted to match the input specifications described in the GEMINI vignette, including (i) a guide-by-sample count matrix, (ii) a replicate annotation file, and (iii) a guide annotation file. Positive and negative control sets were constructed from the simulation labels. For single-gene controls, 50 non-fitness genes (wild-type fitness) were randomly selected as nonessential controls and 50 fitness genes were randomly selected as essential controls. For pairwise negative controls, 50 gene pairs with *Kgi = 1* (no interaction) were randomly sampled from the simulated dataset.

We executed GEMINI (R implementation) using the *gemini_inference* function with the following parameters: *cores = 1, threshold = 0*.*008*, and *n_iterations = 20*. Precision–recall curves were generated using GEMINI “strong scores,” and simulated ground-truth interactions were defined as gene pairs with *Kgi* < 1.

### Comparing other screens: Lenoir et al., Fong et al., Fielden et al. with GRAPE

Raw read count matrices for the combinatorial CRISPR screens from Lenoir et al., Fong et al., and Fielden et al. were obtained from the supplementary data of the respective publications. All datasets were processed using the GRAPE pipeline with study-specific preprocessing steps described below.

The published dataset from Lenoir et al. includes gene–gene pairs as well as constructs paired with control guides. Guide pairs involving control genes were excluded prior to analysis. The remaining dataset was processed directly with GRAPE to compute FCs, predicted additive fitness, and Zgi.

The Fong et al. dataset contains multiple time points for each screened cell line. To ensure consistent screen quality, we performed quality control (QC) by evaluating (i) the separation between essential and nonessential single-gene LFC distributions and (ii) sample clustering across replicates. For each cell line, the time point showing the strongest essential–nonessential separation and highest within-replicate concordance was selected for analysis. The selected samples were: A549 (T24), CAL27 (T22), CAL33 (T17), MDA-MB-231 (T14), MCF7 (T13), and MCF10A (T17). These filtered datasets were subsequently analyzed with GRAPE.

The CRISPRi library used by Fielden et al. contains both perfect-match sgRNAs and single-mismatch variants targeting essential genes. We removed all mismatch-containing constructs and retained only perfect-match guides for downstream analysis. The resulting filtered count matrix was processed with GRAPE following the standard pipeline.

Across all datasets, GRAPE was run with default parameters unless otherwise specified. Predicted fitness, raw GI scores, Zgi, and statistical significance were computed as described above.

### Parameter optimization of SYNULATOR using Bayesian optimization

To generate simulated CRISPR multiplex data that closely recapitulated the empirical distribution of LFC distribution observed in Figure 3F, we optimized SYNULATOR parameters using Bayesian optimization. Simulation parameters were mostly included in the optimization, except the total number of genes (n=167) and the number of guides per gene (n=2) were held constant to match the structure of the library. For each simulation, guide-level FCs were aggregated to the gene level by averaging across guides, ensuring consistency with the real data.

Similarity between simulated and empirical FC distributions was quantified using a composite loss function based on distributional distance metrics. We computed the Wasserstein distance and the two-sample Kolmogorov–Smirnov statistic between real and simulated FC vectors, capturing differences in both global distributional shifts and overall shape. The final loss was defined as a weighted sum of these two metrics and was minimized during optimization.

Bayesian optimization was performed using *Optuna* with a Tree-structured Parzen Estimator (TPE) sampler. Parameter search spaces were defined a priori based on biological plausibility and prior experience with Synulator, and constraints were applied to enforce valid ordering of minimum and maximum fitness and genetic interaction effect sizes.

To account for stochastic variability, each parameter configuration was evaluated using multiple simulation replicates with different random seeds, and the loss was averaged across replicates. Optimization was conducted for a fixed number of trials with parallel execution to improve computational efficiency. The parameter set achieving the minimum loss was selected as the optimal SYNULATOR configuration. The optimal parameters are: *num_total_genes = 167, num_fitness_genes = 53, mu_k_fitness_min=0*.*248, mu_k_fitness_max=0*.*96, genetic_interaction_frequency=0*.*023, genetic_interaction_fitness_min=0*.*21, genetic_interaction_fitness_max=0*.*92, wt_gi_multiplier=0*.*51, num_guides = 2, time=4, transduction_depth=962, median_read_depth = 456, overdispersion_param=0*.*01*.

## Funding

National Institutes of Health/NIGMS grant R35GM130119 (TH) National Institutes of Health/NCI grant R35CA274234 (J Chen) MDACC Odyssey Fellowship Program (CL)

## Author contributions

Conceptualization: JChou, JChen, TH

Methodology: JChou, CL, TH

Investigation: JChou, CL, SK, TH

Visualization: JChou, TH

Supervision: TH, JChen

Writing—original draft: JChou, TH

Writing—review & editing: all authors

## Competing interests

The authors declare no competing interests.

## Data and materials availability

All data supporting the findings of this study are available in the Supplementary Materials. The GRAPE software is publicly available at https://github.com/hart-lab/GRAPE, and the Synulator simulation framework is available at https://github.com/hart-lab/SYNULATOR.

## Notes

https://github.com/hart-lab/GRAPE

https://github.com/hart-lab/SYNULATOR

### Competing Interest Statement

The authors have declared no competing interest.

### Summary of Updates

Added missing author in metadata. Authors are correct in pdf file.

