## Supplementary Information for "Analysis of Combinatorial Knockout CRISPR Screens with GRAPE: Genetic interaction Regression Analysis of Pairwise Effects"

Juihsuan Chou *et al.*

**This PDF file includes:**

Supplementary Text
Figs. S1 to S2
Data S1 to S2

**Supplementary Text**

GRAPE Howto guide

**Overview**

This guide describes how to use GRAPE (Genetic interaction Regression Analysis of Pairwise Effects) to infer genetic interactions from pooled multiplex CRISPR screens. GRAPE supports both symmetric and asymmetric combinatorial library designs and can be applied across different CRISPR modalities.

GRAPE performs three core tasks:

1. Normalization of raw read count data from pooled screens.
2. Regression-based estimation of single-gene fitness effects from dual-perturbation measurements.
3. Compute genetic interaction scores by comparing observed pairwise fitness to predicted additive expectations, followed by variance normalization and statistical testing.

**Input Data Requirements**

1. Read Count file
   - Rows: sgRNA or sgRNA pairs with guide-level identifiers as index and gene-level identifiers as first column.
   - Columns: samples or replicates
   - Values: raw read counts
2. Target Gene file
   A plain text file with one gene per line, includes all unique genes in the library and exclude controls. For asymmetric library designs, combine target genes and query genes into one target gene file.

---

**Using in a Python Notebook:**

**Step 1: Load input read count file and target gene list**

```
import grape as gr

reads = gr.load_readcount_matrix(filepath = 'path_to_read_count_file.txt',
                                 delimiter = '\t')
target_gene_list = gr.load_genelist(filepath ='path_to_target_gene_list.txt')
```

**Parameters**

filepath: The path to the file to be loaded

delimiter: tab or comma delimited file. Defaule is '\t'.

### Step 2: Generate fold change input

#### 2.1 Compute Log2 Fold Change

```
raw_fc = gr.get_foldchange_matrix(reads_df = reads,
                                  control_columns = ['T0_R1', 'T0_R2'])
```

**Parameters**

reads_df : DataFrame containing raw read counts, with index as unique IDs, columns as samples and first column is the target-gene column

control_columns : Either a list of control sample labels (column names) or a list of their column indices

min_reads : optional, minimum read count threshold for filtering samples

pseudocount : optional, small value added to read counts to avoid division by zero and log transformations of zero. Default is 1.

**Returns**

DataFrame with log2 fold-change values, where index is the same as `reads_df` and columns correspond to target samples

#### 2.2 Average fold changes across replicates and guides

```
fc = gr.get_mean_foldchange(fc_df = raw_fc,
                            target_columns = ['T18_R1', 'T18_R2'])
```

**Parameters**

fc_df : DataFrame of log2 fold-change values, where the first column is the target gene(s). Subsequent columns are the replicates to be averaged.

target_columns : List of column labels indicating replicates to be averaged. If None, all columns except the first are used.

no_mean_replicates : bool, optional. Whether to average across replicate columns. Default is False.

no_groupby_targets : bool, optional. Whether to compute the mean fold-change by target gene(s). Default is False.

**Returns**

DataFrame with mean fold-change values, optionally grouped by target genes

#### 2.3 Normalize fold changes

Default: Mode-center normalization. Centers the fold-change distribution so that the mode is zero, assuming most gene pairs have no interaction.

```
modecenter_meanfc = gr.mode_center(mean_fc_df = fc)
```

Optional: reference-gene normalization. Normalizes fold changes so that the median of non-essential genes is centered at zero.

```
modecenter_meanfc = gr.mode_center_vs_reference_genes(mean_fc_df = fc,
                        noness_genes: List[str])
```

**Parameters**

mean_fc_df: DataFrame containing fold-change values, where the index represents target genes

noness_genes: List of non-essential genes

**Returns**

DataFrame of normalized Fold-change values

### Step 3: Fit the GRAPE regression model

#### 3.1 Construct predictor matrix

```
predictor, obs = gr.make_predictor_matrix(
                        fc_df = modecenter_meanfc,
                        target_gene_list = target_gene_list,
                        genepair_del = '_')
```

**Parameters**

fc_df : DataFrame containing normalized fold-change values with index as genes and gene pairs

target_gene_list : List of target genes to be included in the regression model

genepair_del : optional. Delimiter used to separate gene pairs in index names. Default is "_"

**Returns**

- Predictor matrix: binary matrix where columns are target genes and rows are gene constructs
- Response vector: observed fold-change values

#### 3.2 Run regression

```
pairs, singles, model = gr.do_regression(
                              predictor_matrix = predictor,
                              obs_vector = obs,
                              fit_intercept = False,
                              genepair_del = '_')
```

**Parameters**

predictor_matrix : DataFrame of a binary predictor matrix generated from make_predictor_matrix

obs_vector : Response vector generated from make_predictor_matrix
fit_intercept : bool, optional. Whether to fit an intercept in the regression model. Default is False.
genepair_del: optional. Delimiter used to separate gene pairs in index names. Default is "_".

**Returns**

- DataFrame containing observed FC and predicted FC for gene pairs, and single gene FC
- DataFrame containing single-gene FC
- Dictionary with regression metadata (R-squared value, intercept, and model parameters)

### Step 4: Apply dynamic range filter and re-fit regression

#### 4.1 Dynamic range filter

Identify target list where predicted phenotype (fc_exp) is beyond the dynamic range of the assay (min fc_obs). These break the regression model.

```
remove_me = gr.dynamic_range_filter(regression_pairs = pairs)
```

**Parameters**

regression_pairs: DataFrame of 'pairs' output from do_regression()

**Returns**

subset of regression_pairs dataframe whose indices should be deleted from the foldchange df, and run grape do_regression again.

#### 4.2 Remove gene pairs outside dynamic range and re-run regression

```
fc_filt = fc.drop(remove_me.index.values, axis = 0)
modecenter_meanfc_filt = gr.mode_center(mean_fc_df = fc_filt)

predictor, obs = gr.make_predictor_matrix(
                        fc_df = modecenter_meanfc_filt,
                        target_gene_list = target_gene_list,
                        genepair_del = '_')

pairs, singles, model = gr.do_regression(
                              predictor_matrix = predictor,
                              obs_vector = obs,
                              fit_intercept = False,
                              genepair_del = '_')
```

### Step 5: Calculate GI Zscore

```
pairs_localZ = gr.get_zscore(regression_df = pairs,
                             half_window_size = 500,
                             monotone_filter = False)
```

**Parameters**

regression_df : DataFrame containing genetic interaction data, output from `do_regression()`

half_window_size : optional, default=500. Half-window size for calculating local variance. If set to 0, calculates global variance. Half-window size should be adjusted according to the total number of gene pairs in the library, more details in Methods section.

monotone_filter : optional, default=False. If True, ensures a monotonically increasing variance.

**Returns**

Updated DataFrame with added columns:
- `local_std`: Local standard deviation of genetic interactions.
- `GI_Zscore`: Z-score of genetic interactions.
- `Pval_synth`: P-value for synthetic interactions.
- `Padj_synth`: Adjusted p-value (FDR) for synthetic interactions.
- `Pval_supp`: P-value for suppressing interactions.
- `Padj_supp`: Adjusted p-value (FDR) for suppressing interactions.

---

**Command-Line Usage**

Download and installation instructions are provided on the GRAPE GitHub repository: https://github.com/hart-lab/GRAPE

GRAPE can be run directly from the command line:

```
$bash

grape \
  -i path/to/read_count_file.csv \
  -o output_directory/ \
  -c T0_R1 T0_R2 \
  -t path/to/target_gene_list.txt \
  --target-columns T18_R1 T18_R2 \
  -p T18
```

Run `grape --help` for all available options.

Synulator Howto guide

**Overview**

This guide covers the two main steps for using synulator to simulate genetic interaction screen data.

Synulator simulates CRISPR-based genetic interaction screens by:

1. Generating a fitness matrix representing single knockouts and pairwise interactions. The fitness matrix represents contains the "ground truth" of single and double knockout fitness and genetic interaction fitness effect. The fitness matrix can either be generated randomly using this function or manually generated by the user.

2. Converting that matrix into a fold change table with realistic experimental noise. The fold change table can then be used to evaluate actual experimental data

---

**Using in a Python Notebook:**

**Step 1: Generate a fitness matrix**

The fitness matrix is an N×N matrix of *k* values where:

- **Diagonal entries** represent single gene knockout fitness ($k_{gene}$ = 1.0 for wildtype, 0.0 for lethal)
- **Off-diagonal entries** represent genetic interactions between gene pairs ($k_{gi}$ = 1.0 for no interaction, 0.0 for synthetic lethal)
- The actual fitness, k, of a dual knockout of $gene_i$ and $gene_j$ is therefore $(k_i)(k_j)(k_{gi})$

```
from synulator import generate_fitness_matrix

fitness_matrix, gene_labels = generate_fitness_matrix(
    num_total_genes = 200,            # Total genes in library
    num_fitness_genes = 50,           # Genes with fitness defects when KO
    mu_k_wt = 1.0,                    # Fitness value for wildtype knockouts
    mu_k_fitness_min = 0.2,           # Min fitness for fitness genes
    mu_k_fitness_max = 1.0,           # Max fitness for fitness genes
    genetic_interaction_frequency = 0.01,      # Probability of interaction
    genetic_interaction_fitness_min = 0.2,     # Min interaction fitness
    genetic_interaction_fitness_max = 1.0,     # Max interaction fitness
    wt_gi_multiplier=0.1              # Reduces GI frequency for WT-WT pairs
)
```

**Parameters**

num_total_genes: the number of genes in your synthetic all-by-all genetic interaction screen.

num_fitness_genes: the number of genes, out of the total, that have a single knockout fitness defect (including 'essential' genes). All other genes are assumed to have k=1.

mu_k_wt: fitness for wildtype genes; i.e. knockout fitness. Default is 1.0 and should not be changed under normal conditions.

mu_k_fitness_min, max: the range of fitness values for the fitness genes. For each fitness gene, k is drawn randomly from Uniform(min, max).

genetic_interaction_frequency: the probability for any gene pair to have a genetic interaction. Range for 0.01-0.10 is reasonable.

wt_gi_multiplier: a number less than 1.0 to reduce the frequency of genetic interactions between wildtype genes. In general, there are more candidate interactions between wildtype genes than between gene pairs including at least one fitness gene. This can result in an excess of synthetic genetic interactions with expected ($k_i k_j$, non-interaction) fitness of 1.0, which is rare in real experiments. This correction factor spreads out the interactions across the range of $k_i k_j$ expected fitness values.

genetic_interaction_fitness_min, max: the range of fitness values for genetic interactions. If a gene pair is randomly assigned a genetic interaction, the kgi is drawn from U(min,max).

Example: for the listed parameters above, a 200x200 fitness matrix will be generated. For the 200 synthetic genes, 50 are “fitness genes” with ki drawn from U(0.2,1.0). The remaining 150 are assigned ki=1.0. These single gene fitness values are the diagonal of the matrix.

Then, for each gene pair (off-diagonal), $k_{gi}$ is determined. Whether a genetic interaction is present is determined by the genetic interaction frequency and, if applicable, the wt_gi_multiplier. If a genetic interaction is present, $k_{gi}$ is drawn from U(genetic interaction fitness min, genetic interaction fitness max); otherwise $k_{gi}$ = 1.0.

The result is a 200x200 matrix with single knockout fitness on the diagonal (ki). Expected double knockout fitness can be calculated from ($k_i k_j$), and true double knockout fitness is calculated from ($k_i k_j k_{ij}$), where $k_{ij}$ is the genetic interaction fitness/effect size $k_{gi}$.

**Output**

- `fitness_matrix`: pandas DataFrame (N×N) with fitness and genetic interaction values
- `gene_labels`: Dictionary categorizing genes as 'wildtype' or 'fitness'

**<u>Step 2: Generate a fold change table</u>**

In this step, data from a fitness matrix is used to construct a synthetic population growth experiment. A sample is drawn from the endpoint and compared to the start to generate a fold change table. Experimental and technical noise is added to maximize similarity with real-world CRISPR screening data. The fitness matrix can be generated using the SYNULATOR method above or constructed manually for those desiring more granular control over the simulations.

```
from synulator import generate_fitness_table

fitness_table = generate_fitness_table(
    fitness_matrix = fitness_matrix,    # From Step 1
    gene_labels = gene_labels,          # From Step 1
    num_guides = 4,                     # gRNAs per target
    sigma_k = 0.03,                     # Noise in fitness measurements
    time = 8,                           # Cell doublings
    transduction_depth = 500,           # Initial cell count
    median_read_depth = 500,            # Sequencing depth
    overdispersion_param = 0.5,         # Negative binomial noise (0-1)
    pseudocount = 1,                    # Prevents zero read counts
    seed = 42                           # For reproducibility
)
```

**Parameters**

fitness_matrix: a pandas dataframe containing a symmetric fitness matrix.

gene_labels: a dictionary categorizing synthetic genes in the fitness matrix as 'wildtype' or 'fitness'

num_guides: the number of gRNA targeting each gene/gene pair. This is included only to make output compatible with downstream bioinformatic pipelines that require multiple reagents per gene target. An idealized fold change table is generated with num_guides=1. For num_guides > 1, individual guides have effect sizes normally distributed around the target effect size; i.e. $k_i k_j k_{ij.}$

sigma_k: Sigma k ($\sigma_k$) is the standard deviation of the integrated noise term. The *true* fitness of a gene (or gene pair) is $k_i$ (or $k_i k_j k_{ij}$). The *observed* growth rate of cells is:

$$X_t = X_0 * 2^{kt + \varepsilon}, \text{ where } \varepsilon \sim N(0,\sigma_k).$$

This value is empirically estimated; reasonable values range from 0.02-0.05. In the absence of this term, the observed fold change for all wildtype genes ($k_i$=1.0) is exactly zero. With this term, fold changes for wildtype genes are approximately normally distributed with standard deviation $\sigma_k t$, where t is the number of cell doublings.

time: number of doublings. See t term above.

transduction_depth: initial cell count per gene/gene pair target at to. This is the X0 term in the above growth equation.

median_read_depth: this is the "sampling depth" for targets at the Xt timepoint. Higher read depth implies larger counts, and therefore lower variability, at high fold change. Most experiments are 200x-1000x median read depth.

overdispersion_param: Endpoint cells, Xt, are sampled at median_read_depth and then binomial noise is added using this parameter. This mimics with high fidelity the increased variability of CRISPR targets with severe fitness defects.

seed: randomization seed. Set the seed for exact replicates or use without seed to generate random fold change tables from the same fitness table (i.e. technical replicates).

#### Output Columns

| Column | Description |
|---|---|
| `guide_id` | Unique guide identifier |
| `target_id` | Target gene(s) |
| `mu_k` | True fitness value |
| `GI` | True genetic interaction term |
| `label` | Gene type classification |
| `X0` | Initial cell count |
| `k_obs` | Observed fitness value |
| `Xt` | Final cell count |
| `Reads_t` | Simulated read counts |
| `Log2fc` | Log2 fold change |

---

#### Command-Line Usage

Download and installation instructions are provided on the SYNULATOR GitHub repository: https://github.com/hart-lab/SYNULATOR

Both steps can be run together via the CLI:

```
$bash
synulator -o simulation_results.txt \
    --num_total_genes 300 \
    --num_fitness_genes 50 \
    --time 6 \
    --seed 42
```

Run `synulator --help` for all available options.

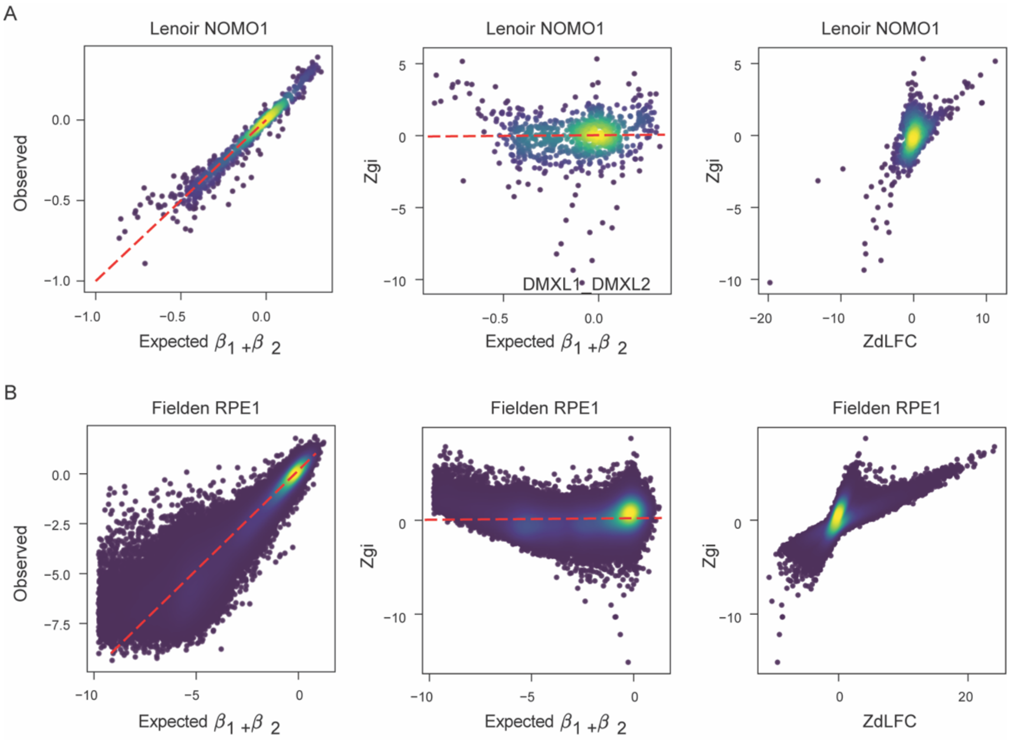


**Fig. S1. GRAPE analysis across 2 combinatorial CRISPR datasets.**

Scatter plots showing expected FC versus observed FC, expected FC versus Zgi, and ZdLFC versus Zgi derived from GRAPE analysis for **(A)** the NOMO1 cell line from Lenoir et al. and **(B)** the RPE1 cell line from Fielden et al.

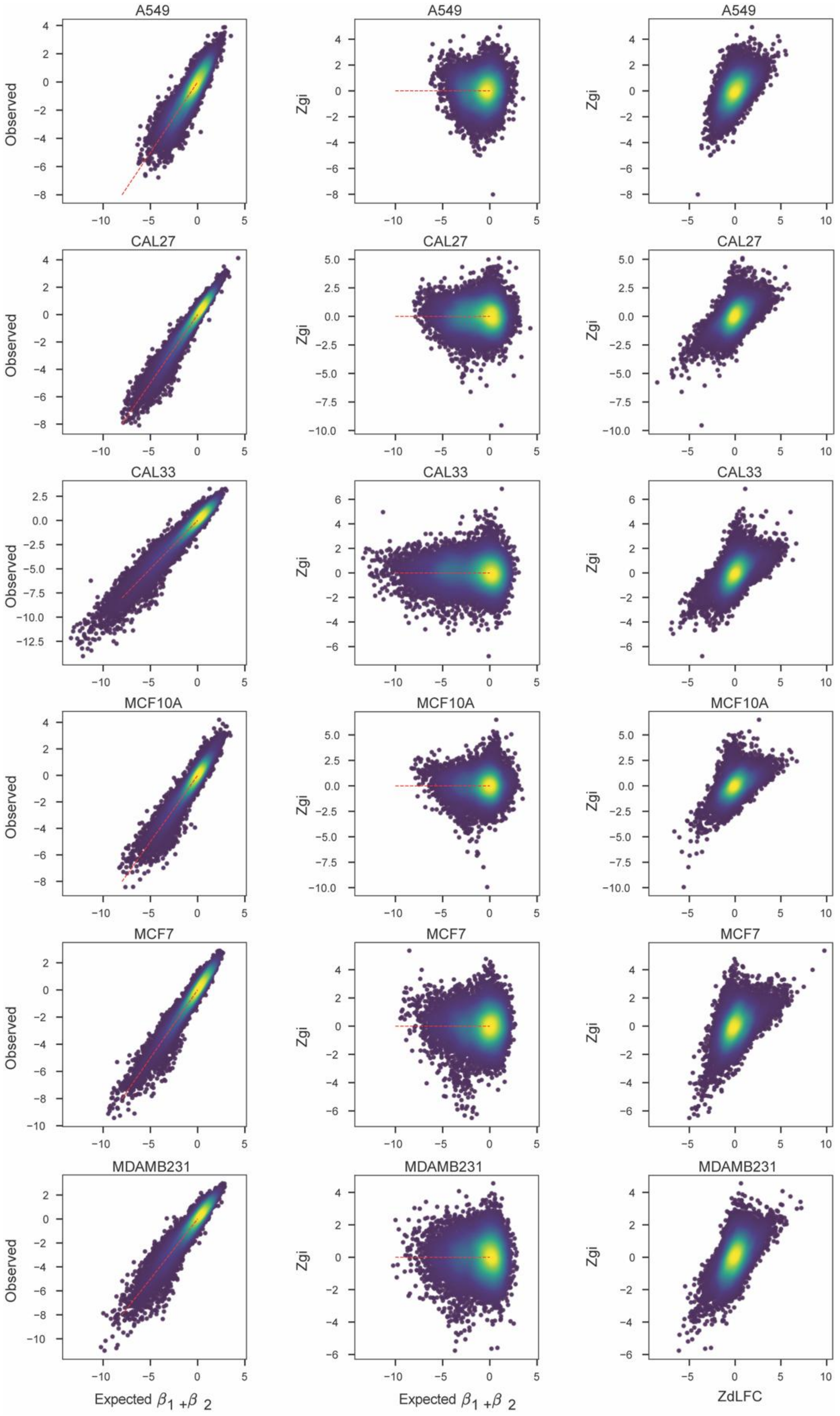


**Fig. S2. GRAPE analysis across multiple cell lines from Fong et al.**

Scatter plots showing expected FC versus observed FC, expected FC versus Zgi, and ZdLFC versus Zgi for six cell lines analyzed in Fong et al.

**Data S1. (separate file)**
GRAPE output for the PC9 cell line with DDR library.

**Data S2. (separate file)**
GRAPE outputs for 9 published combinatorial CRISPR screens.
